# Functional and evolutionary significance of unknown genes from uncultivated taxa

**DOI:** 10.1101/2022.01.26.477801

**Authors:** Álvaro Rodríguez del Río, Joaquín Giner-Lamia, Carlos P. Cantalapiedra, Jorge Botas, Ziqi Deng, Ana Hernández-Plaza, Lucas Paoli, Thomas S.B. Schmidt, Shinichi Sunagawa, Peer Bork, Luis Pedro Coelho, Jaime Huerta-Cepas

**Affiliations:** Centro de Biotecnología y Genómica de Plantas, Universidad Politécnica de Madrid (UPM) - Instituto Nacional de Investigación y Tecnología Agraria y Alimentaria (INIA-CSIC), Campus de Montegancedo-UPM, 28223, Madrid, Spain; Departamento de Biotecnología-Biología Vegetal, Escuela Técnica Superior de Ingeniería Agronómica, Alimentaria y de Biosistemas, Universidad Politécnica de Madrid (UPM), Madrid, Spain; Department of Biology, Institute of Microbiology and Swiss Institute of Bioinformatics, ETH Zürich, Zürich 8093, Switzerland; Structural and Computational Biology Unit, European Molecular Biology Laboratory, D-69117 Heidelberg, Germany; Max Delbrück Centre for Molecular Medicine, Berlin, Germany; Yonsei Frontier Lab (YFL), Yonsei University, Seoul, South Korea; Department of Bioinformatics, Biocenter, University of Würzburg, Würzburg, Germany; Institute of Science and Technology for Brain-Inspired Intelligence, Fudan University, Shanghai, China; MOE Key Laboratory of Computational Neuroscience and Brain-Inspired Intelligence, and MOE Frontiers Center for Brain Science, Shanghai, China

## Abstract

Most microbes on our planet remain uncultured and poorly studied. Recent efforts to catalog their genetic diversity have revealed that a significant fraction of the observed microbial genes are functional and evolutionary untraceable, lacking homologs in reference databases. Despite their potential biological value, these apparently unrelated orphan genes from uncultivated taxa have been routinely discarded in metagenomics surveys. Here, we analyzed a global multi-habitat dataset covering 151,697 medium and high-quality metagenome assembled genomes (MAGs), 5,969 single-amplified genomes (SAGs), and 19,642 reference genomes, and identified 413,335 highly curated novel protein families under strong purifying selection out of previously considered orphan genes. These new protein families, representing a three-fold increase over the total number of prokaryotic orthologous groups described to date, spread out across the prokaryote phylogeny, can span multiple habitats, and are notably overrepresented in recently discovered taxa. By genomic context analysis, we pinpointed thousands of unknown protein families to phylogenetically conserved operons linked to energy production, xenobiotic metabolism and microbial resistance. Most remarkably, we found 980 previously neglected protein families that can accurately distinguish entire uncultivated phyla, classes, and orders, likely representing synapomorphic traits that fostered their divergence. The systematic curation and evolutionary analysis of the unique genetic repertoire of uncultivated taxa opens new avenues for understanding the biology and ecological roles of poorly explored lineages at a global scale.

## Introduction

Over the last decades, the sequencing of environmental DNA by metagenomics surveys has unveiled a great level of microbial biodiversity. These efforts have not only led to the discovery of new bacterial and archaeal lineages, most of which have not yet been isolated and cultured, but also uncovered an enormous amount of unknown prokaryotic genes whose significance is still largely unexplored.

Recent large scale microbiome studies covering human gut^1^, ocean^2^ and multi-habitat samples^3^, consistently report that a large proportion (27%–47%) of the observed metagenomic genes are functional and evolutionary untraceable (i.e., no homologs in reference databases). The lack of both functional information and evolutionary links to known organisms makes these supposedly orphan genes difficult to interpret, curate, and integrate into comparative metagenomic pipelines, being usually discarded from most analysis. As a result, the unique genetic repertoire of uncultivated microbial lineages has not yet been incorporated into reference databases of protein domains, orthologous groups (OGs) or microbial gene families^4^. Thus, despite the astonishing amount of data generated by metagenomics sequencing, their study still rely on reference resources that are heavily biased towards the genetic pool of fully sequenced microorganisms, leaving a gap in our knowledge base and impeding our ability to investigate the true diversity of microbially encoded genes on Earth^5,6^.

The distinctive genetic repertoire of uncultivated bacteria and archaea might be key to understanding the biology and evolution of new microbial lineages^7,8^. Recent examples of novel molecular functions discovered out of unknown genes from uncultivated taxa include new enzymes^9,10^, antibiotics^11^, and thousands of putatively functional small peptides^12^. Similarly, the cultivation and genomic analysis of poorly explored organisms (e.g., Asgard archaea^13^ or Planctomycetes^8^) has led to the discovery of new metabolisms and unusual biology^14^. However, at a global scale, our current view of the genetic repertoire of uncultivated organisms remains anecdotal. Not only the function of many of their genes is ignored, but also their potential evolutionary links with other uncultured species, their selective pressures and ecological significance.

Here, we argue that recent methodological advances have unlocked most important impediments at the technical and computational level to perform de novo comparative genomics analyses of uncultivated taxa at a global scale. Firstly, hundreds of high-quality metagenome-assembled genomes are publicly available^1,3,15^ and can be taxonomically classified^16^. Secondly, it is now possible to use sensitive homology sequence searches and clustering algorithms on huge genomic catalogs^17,18^. This has allowed recent metagenomic surveys to employ the concept of protein families — rather than individual gene entries^19^ — to identify and quantify genetic novelty^15,20^, contributing to the annotation of distant homologs and reducing the number of putative orphan genes. Most importantly, these advances have laid the foundation for the systematic analysis of thousands of uncultured organisms under a common phylogenomic framework, which allowed us to address long-standing questions on the extent and biological value of their seemingly unrelated orphan genes.

## Results

### A curated catalog of unknown protein families specific from uncultivated taxa

We used a comprehensive comparative genomics approach to identify novel protein families in a large multi-habitat metagenomic dataset covering 151,697 medium and high-quality metagenome assembled genomes (MAGs), 5,969 single-amplified genomes (SAGs), and 19,642 reference genomes (i.e., from fully sequenced species). This set includes over 400 million gene predictions and was assembled by unifying five data sources spanning 82 habitats: two MAG collection spanning thousands of samples from diverse origins (GEM^3^ and GMGC^15^), a comprehensive human gut catalog (UHGG)^1^, a global ocean catalog (OMD)^21^, and the GTDB r95 reference database^22^ (Table S1).

Considering that the large pool of unknown metagenomic sequences is typically enriched in incomplete genes, assembling artifacts, distant homologs and potential pseudogenes, we applied strict quality and novelty filters to our protein family discovery approach (summarized in Fig. 1A), aiming at compiling a collection of high-quality novel protein families from uncultivated taxa. First, we used deep homology clustering at the amino acid level to group all genes into sequence clusters, each serving as a proxy for a different protein family. Next, to ensure the detection of unknown protein families specific of uncultivated species, we selected clusters lacking sequence members belonging to any of the known reference genomes included in our dataset, and excluded entries with significant hits against the latest versions of PFAM-A/B^23^, eggNOG^24^, and RefSeq^25^. This allowed us to identify clusters composed exclusively of proteins without known homologs. Moreover, to ensure the identification of functionally and evolutionarily relevant protein families, we only selected protein clusters containing at least three sequences from three different uncultivated species, and with a conserved aligned region of at least 20 contiguous amino acids. This allowed us to discard clusters inferred from barely overlapping sequence segments, as well as to build a unique genomic signature for each protein family (i.e., putative novel protein domains). To avoid pseudogene-based and viral-specific protein families, we also excluded clusters matching either the AntiFam^26^ or pVOGs^27^ databases. Finally, we required genes covered by the selected novel families to be under purifying selection (dN/dS < 0.5, Fig. 1B) as expected for functional coding sequences^28^, and to be expressible, either based on *in silico* prediction methods or by empirical evidence (i.e., with significant hits against recent metatranscriptome surveys^2,29^).

**Figure 1.**
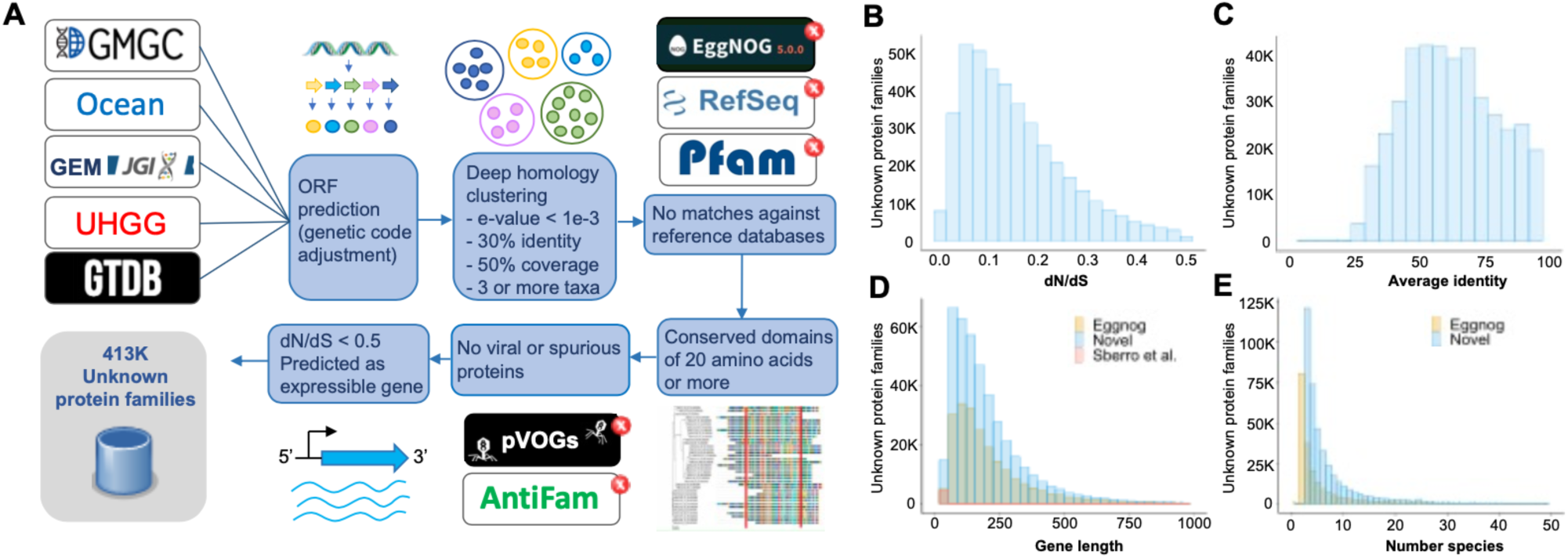
Protein family discovery pipeline and general statistics. **(A)** Workflow used to identify high quality unknown protein families. **(B)** dN/dS distribution of the unknown protein families. **(C)** Average protein identity of the unknown protein families. **(D)** Protein length distribution of unknown families (blue histogram) compared to bacterial and archaeal eggNOG orthologous groups (yellow) and novel small peptides reported in Sberro et al.^12^ (red) **(E)** Number of species distribution of unknown families (blue) compared to bacterial and archaeal eggNOG orthologous groups (yellow).

Our final catalog includes a total of 413,335 previously unknown and highly conserved protein families, with size, length, and species-content distributions comparable to those currently found in global microbial databases (Fig. 1D and 1E). This constitutes an almost three-fold increase over the total number of prokaryotic OGs described to date (namely 219,934 bacterial and archaeal eggNOG OGs), evidencing the high number of putative molecular functions that are commonly neglected in current metagenomic surveys (Fig. 1D)^30,31^. From an evolutionary perspective, the identified protein families are conserved (average identity 63.2%, Fig. S1) and all qualify as novel orthologous groups at the bacterial or archaeal level using phylogeny-based orthology prediction methods. Compared to global microbial databases, we found that unknown protein families are slightly enriched in transmembrane and signal peptide-containing proteins (being 7.6% and 7.9% more frequent than in eggNOG, respectively), which suggests that they may play an important role in mediating interactions with the environment. Moreover, while we did not specifically target small peptides, we identified 13,456 families of proteins shorter than 50 residues, 486 of which have been reported previously as novel functional genes^12^, 257 containing antimicrobial signatures (Table S2). The lack of distant paralogs within the inferred protein clusters, together with the strong purifying selection measurements (average dN/dS of 0.15), reinforces the idea of each cluster representing an unknown but highly conserved molecular function. In order to facilitate the exploration of this large catalog and its integration into future studies, we provide an online interactive browser and extensive downloading material at http://novelfams.cgmlab.org.

### Conserved synteny between novel protein families and metabolic and resistance genes

Intrigued by the biological significance of the identified unknown protein families, we investigated their phylogenetic, functional, and ecological significance. Firstly, we inferred their putative functional roles by reconstructing their genomic neighborhood and estimating their degree of gene order and functional context conservation, which is a method commonly used for prokaryotic genome analysis^32^. Our results yielded 74,356 (17.98%) novel protein families in phylogenetically conserved operon regions containing at least one contiguous functionally annotated gene in the same DNA strand. Of those, 1,344 families share a genomic context with known and highly conserved marker genes related to energy production or xenobiotic compound degradation pathways (Fig. 2A, Table S3), indicating that the role of unknown protein families may not be limited to accessory molecular functions but could also involve central metabolic processes. As an example, we found 5 novel protein families embedded in operonic regions associated with 3 major pathways of the nitrogen cycle (Fig. 2B). Additionally, we found 965 unknown protein families in the genomic context of well-known antibiotic resistance genes, 25 of which are embedded in clear genomic islands with more than 3 resistance-related neighbor genes (Fig. 2C, Table S4). Moreover, we mapped the protein family signatures derived from our catalog against the set of 11,779 unknown genes recently annotated based on genome-wide mutant fitness experiments^33^, and found 69 matches to genes associated with specific growth conditions (Table S5).

**Figure 2.**
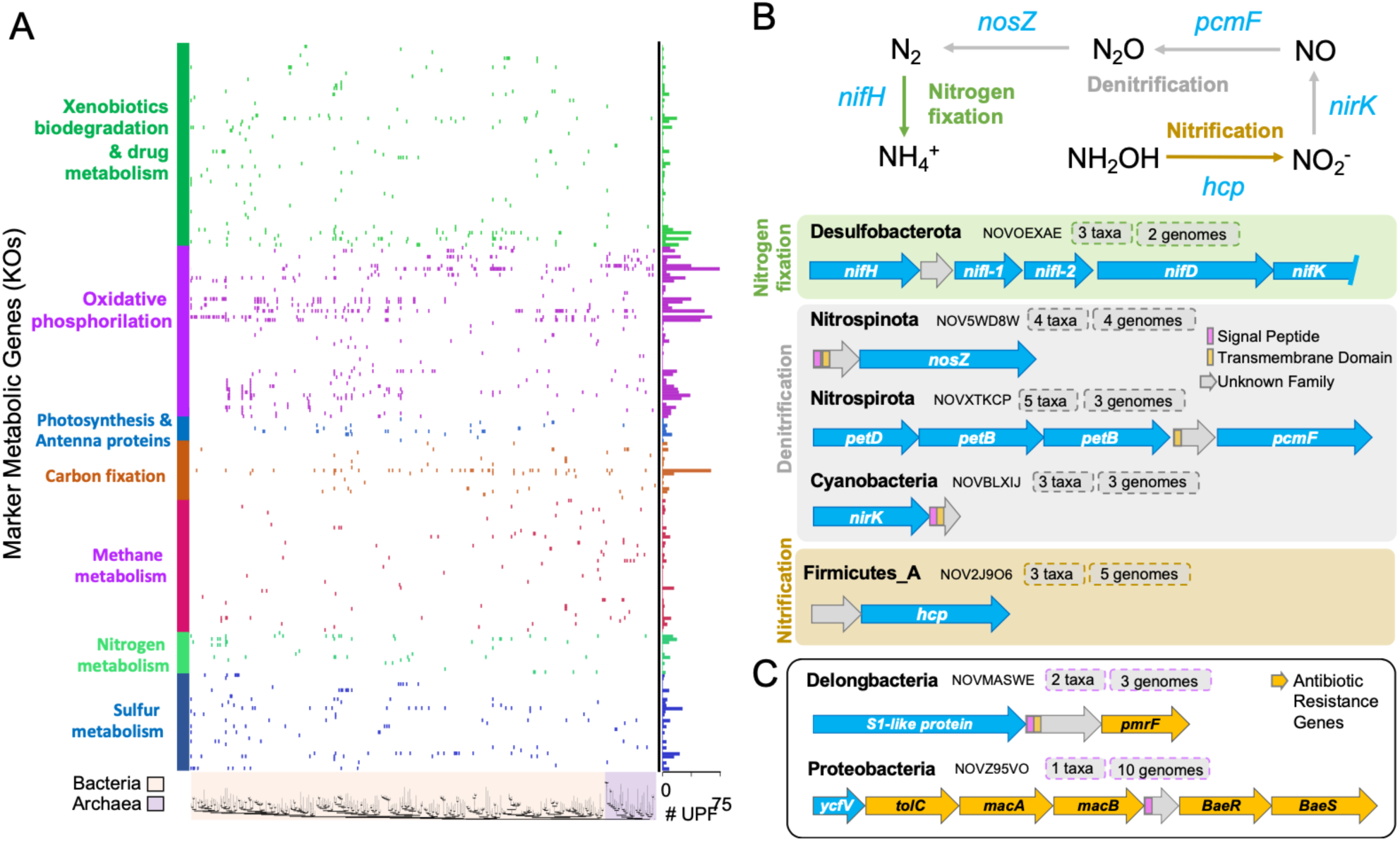
Summary of unknown protein families linked to metabolic marker genes. **(A)** Presence/absence matrix of unknown protein families forming operon-like structures with marker genes involved in energy and xenobiotic degradation KEGG^39^ pathways (rows) across the bacterial and archaeal GTDB phylogeny collapsed at the order level (columns). Taxonomic orders without detections are not shown. The novel protein families associated with each KEGG module can be explored at http://novelfams.cgmlab.org **(B)** Examples of unknown protein families tightly coupled with genes for every nitrogen cycling step. **(C)** Examples of unknown families surrounded by antibiotic resistance genes (as predicted by CARD).

### High content of unknown protein families in the genomes of uncultivated taxa

From a taxonomic point of view, our results reveal that novel protein families are broadly distributed across the entire microbial phylogeny (Fig. 3A), comprising an important fraction of the genetic repertoire of uncultivated taxa. For instance, recently discovered lineages such as Obscuribacterales (a non-photosynthetic sister group to Cyanobacteria^34^) and Thorarchaeales (sulfur-reducing Asgard archaea^35^) contain an average of 616 and 375 unknown protein families from our catalog, per genome, respectively. Even the reduced-size genomes from the ubiquitous and host-dependent Patescibacteria phylum contain a total of 29,935 novel protein families, 63 per genome on average. The majority of these novel families (56%) carry either transmembrane domains or signal peptides, being likely involved in cell-cell or cell-environment interactions such as surface attachment to their hosts and the uptake of compounds^36,37^. Functional predictions based on strict synteny analysis support this idea, with 502 novel families from the Patescibacteria group potentially involved in molecular transportation, 34 in adhesion, and 13 in cytokinesis. Specific examples include the Patescibacteria protein families NOVF69IJ, NOVT9VCK and NOVO6T3T, which were found embedded in genomic contexts clearly associated with hydrolysis, competence and endonuclease degradation; all of them processes related with the disruption of the host cell wall and the incorporation and degradation of DNA^38^.

**Figure 3.**
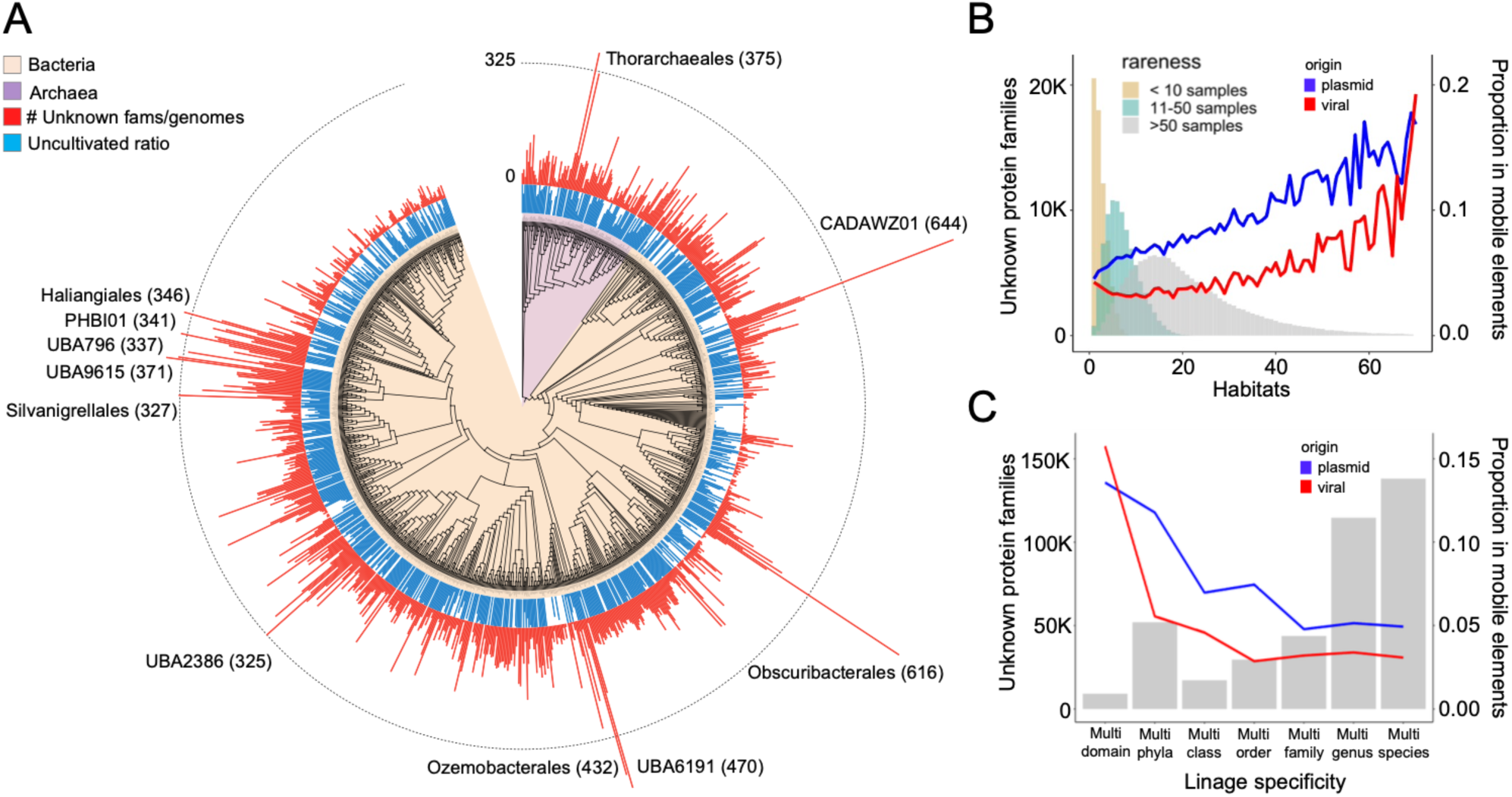
Novel protein families are widespread across the microbial phylogeny and cross habitats. **(A)** Phylogenetic distribution of unknown protein families across the GTDB^22^ bacterial and archaeal phylogeny collapsed at the order level. Red bars indicate the number of unknown protein families per genome in each taxonomic order. Blue bars represent the proportion of uncultivated species under each collapsed order. Branches with more than 400 unknown protein families are indicated. **(B)** Ecological breadth (measured as the number of habitats) of the unknown protein families classified by three levels of rareness (number of samples in which they are detected). The blue and red lines indicate the proportion of protein families predicted as mobile in plasmids and viral contigs respectively, which correlates with ecological breadth. **(C)** Number of unknown protein families confined to each taxonomic rank. The term genus in the x-axis indicates the number of protein families detected in multiple species from the same genus, while the domain bar indicates families spanning more than one phylum from the same domain. The blue and red lines indicate the proportion of protein families predicted as mobile in plasmids and viral contigs respectively.

### Unknown protein families are prevalent across samples and habitats

In addition to their potential functional relevance, we also investigated whether unknown protein families represent rare or widespread traits at a global ecological scale. For this purpose, we estimated the ecological distribution of each protein family in our catalog by mapping their genomic signatures against an expanded set of 63,410 public metagenomic samples spanning 157 habitats (Table S6). Strikingly, we found that the majority of the new protein families (55%) are detected in more than ten samples, span at least two habitats, and are found in microbial communities with very different structures (average Bray-Curtis dissimilarity among samples containing each protein family is 0.88, Fig. S2). This result contrasts with the habitat-specific pattern observed for the majority of individual species-level genes^15^ and indicates that the protein families reported here either represent putative core molecular functions from widespread microbial lineages, or derive from promiscuous mobile elements. To quantify the contribution of each scenario, we specifically identified protein clusters containing at least one sequence member in a plasmid-like or viral-like contig and studied their ecological pattern. Indeed, we found that mobility strongly correlates with the ecological prevalence of the protein families (blue and red lines in Fig. 3B, Fig. S3), although cross-habitat protein families are not restricted to those linked to putative horizontal transfers. The same pattern was observed when analyzing the taxonomic breadth of each individual novel protein family: we found that protein families detected across different high-level taxonomic lineages (e.g., cross-domain and cross-phylum) are more likely to be part of mobile elements (Fig 3C). However, the vast majority of families (90.5%) appeared to be unrelated to obvious events of horizontal gene transfer, suggesting a more constitutive role in their host genomes.

### Synapomorphic protein families of uncultivated phyla, classes and orders

Although all protein families in our catalog could be considered relevant from an evolutionary perspective (e.g., are phylogenetically conserved and under purifying selection), we identified a core set of 980 protein family clusters synapomorphic for entire uncultivated lineages —that is, present in nearly all MAGs/SAGs from a given lineage (90% coverage) but never detected in other taxa. While a similar pattern might be expected for accessory functions at narrow phylogenetic ranges (i.e., species- or genus-specific proteins), these newly discovered protein families can accurately distinguish 16 uncultivated phyla, 19 classes, and 90 orders, involving 179, 104, and 697 novel protein families, respectively. We argue that, despite being unknown and originally treated as orphan sequences, these synapomorphic protein families carry great evolutionary significance and might represent functional innovations that fostered the ancestral divergence and selection of the underlying lineages. Consistent with this idea, synapomorphic protein families show a significantly lower dN/dS ratio than other conserved (but not synapomorphic) unknown protein clusters (Fig. 4B), indicating stronger purifying selection. Similarly, the rate of unknown synapomorphic protein families was higher on uncultivated lineages with poorly understood biology. This is the case of the recently proposed Riflebacteria phylum (25 putative synapomorphic detections) or the Thorarchaeia class, where the 12 detected putative synapomorphic families may help to understand their radiation from other Asgard. The genomic context of these crucial families allowed us to hypothesize about their possible functional role and biological relevance. For instance, we found the Thorarchaeia synapomorphic family NOV4IF0P to be embedded in a highly-conserved genomic context containing eukaryotic-like genes related to protein translation (*rpi35a* and *elp3*; Figure 4C). Similarly, synapomorphic protein families in other lineages also relate to key processes, probably representing functional innovations accounting for their divergence and radiation. For instance, we found synapomorphic families in genomic contexts involved in cytochrome type-c biogenesis (*ccmC, ccmE* and *ccmF*) in the HRBIN17 class, DNA repair (*priA* and *dskA*) in the UBP6 phylum, or chemotaxis (*mcp* and *dgt*) in the Riflebacteria phylum (Figure 4C), among many others that can be easily explored in our online resource.

**Figure 4.**
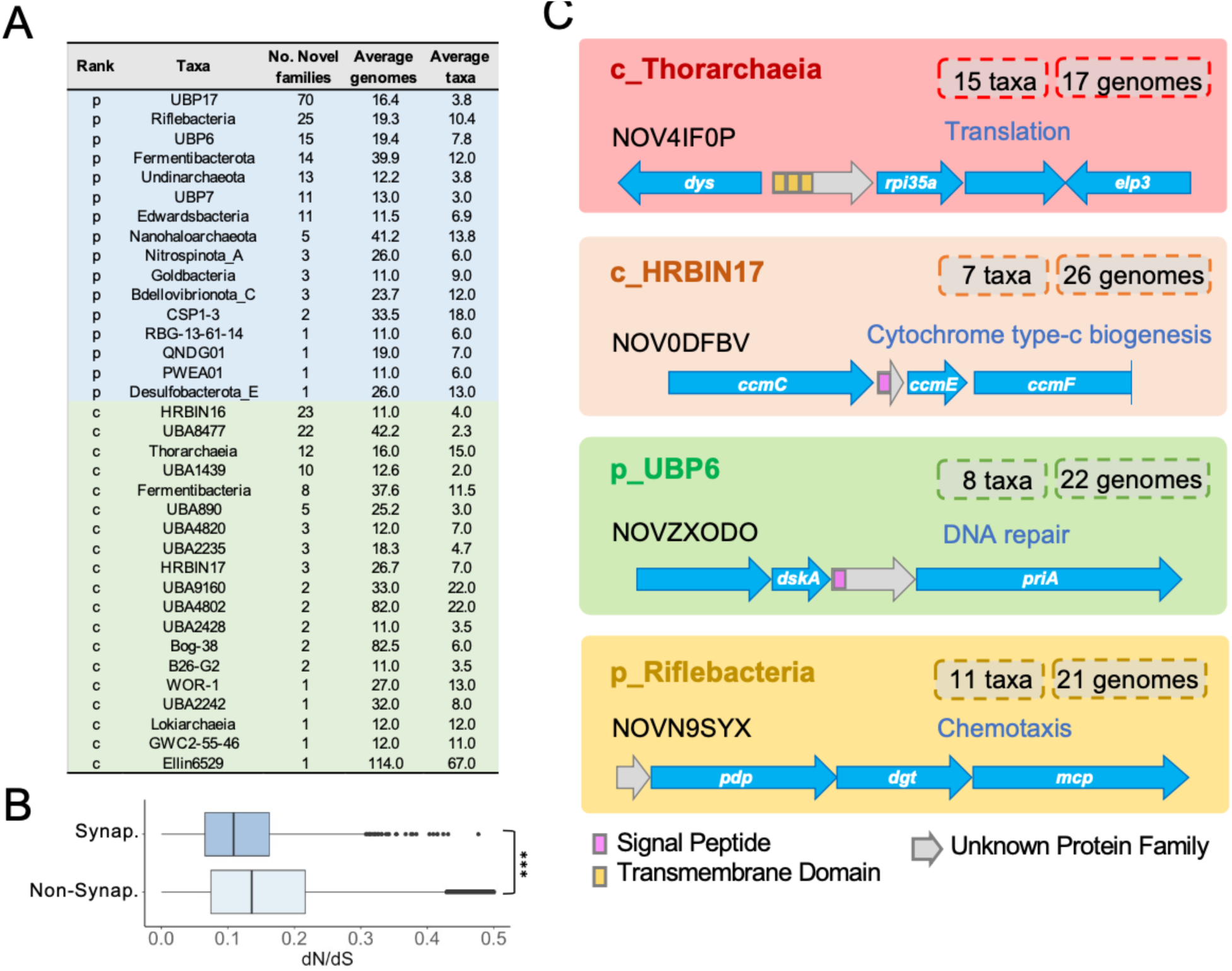
Sample of the 980 unknown protein families highly exclusive and widely distributed (synapomorphic) across different phyla and classes. **(A)** Number of synapomorphic unknown protein families found for 16 uncultivated phyla and 19 classes. Individual examples and other synapomorphic protein families can be explored at http://novelfams.cgmlab.org **(B)** Comparison of dN/dS values between synapomorphic and non-synapomorphic families (Wilcoxon test, two-sided, p-value < 2.2e-16) **(C)** Schematic overview of four phylum and class-level synapomorphic protein families located in conserved gene clusters involved in relevant cellular processes. Gene names correspond to: dys, deoxyhypusine synthase; rpl35ae, large subunit ribosomal protein L35Ae; elp3, Elongation Protein 3 Homolog; rbsK, ribokinase; BET3, trafficking protein particle complex subunit 3; ccmC, heme exporter protein C; ccmE, cytochrome c-type biogenesis protein; ccmF, cytochrome c-type biogenesis protein; dksA, DnaK suppressor protein; priA, primosomal protein N’; pdp, pyrimidine-nucleoside phosphorylase; dgt, dGTPase; mcp, methyl-accepting chemotaxis protein.

## Discussion

Overall, our work provides a global phylogenomic analysis of the largely uncharted repertoire of unique genes from uncultured prokaryotes, serving as a base resource for further investigations on their functional and ecological roles. We demonstrate that evolutionary untraceable sequences from metagenomic surveys are not necessarily orphan genes, pseudogenes or sequencing artifacts. Often, they can represent highly-conserved protein families never observed in cultured organisms but prevalent across unknown lineages and under strong purifying selection. By using strict quality filters, we provide a curated set of relevant novel protein families likely accounting for the diversification of complete high-rank lineages and/or involved in central processes such as energy production, xenobiotic metabolism and microbial resistance. Here, we urge for their incorporation into reference databases and future metagenomic workflows, as they might be key for understanding the biology of poorly studied bacteria and archaeal groups.

Moreover, our work describes a general framework for the curation of unknown metagenomics sequences using phylogenomic techniques, demonstrating that current computational resources are now sufficient to attempt the analysis of unknown metagenomic sequences from a comprehensive comparative genomics perspective. However, based on the large number of genes and protein families that we discarded due to quality filters and limited sampling depth, the data reported here could only represent the tip of the iceberg. We expect that the number of novel protein families inferred from uncultivated organisms using similar frameworks will increase exponentially as we advance in the reconstruction of eukaryotic MAGs and the exploration of the rare biosphere in understudied microbial ecosystems.

## Materials and Methods

### Data collection

We retrieved medium and high quality Metagenome Assembled Genomes (MAGs), Single-amplified Genomes (SAGs) and reference genomes from 5 different studies: i) 51,634 medium and high-quality MAGs (>=50% complete, <=5% contamination) from a planetary multi habitat catalog including >10k samples and covering diverse habitats (GEM)^3^, ii) 46,655 high-quality MAGs (>=90% complete, <=5% contamination) released by another multi habitat catalog spanning 13,175 samples and 14 biomes (GMGC)^15^, iii) 31,910 MAGs from the GTDB-r95 database^40^ iv) 27,123 medium and high-quality MAGs (>=50% complete, <10% contamination), 5,969 Single Amplified Genomes (SAGs) and 1,707 reference genomes obtained from ocean samples^21,41–43^ and v) 4,644 medium and high quality reference human gut MAGs (>=50% complete, <5% contamination) from the UHGG human gut catalog^1^.

### Recalling Open Reading Frames

We observed that some MAG collections contained genes predicted under an incorrect codon table (for instance, genes from the Gracilibacteria or Mycoplasma lineages were predicted under the standard codon table). We therefore re-computed ORF predictions for the MAGs in the GEM, GMGC and GTDB catalogs. For each MAG, we used PROKKA^44^ selecting the correct genetic table for each genome based on its taxonomic annotation. We further verified that the corrected Gracilis Bacteria and Mycoplasma ORFs were indeed longer than the original ones. We obtained 116,208,548, 110,913,525 and 106,052,079 genes for GEM, GMGC and GTDB respectively. After combining them with the 56,637,438 and 10,002,521 genes in the oceanic and UHGG genomes, MAGs and SAGs, we obtained a final catalog of 399,814,111 genes.

### Taxonomic annotations

In order to obtain homogeneous taxonomic annotations for all the MAGs in our collection, we re-annotated them using GTDB-Tk v1.6.0 (GTDB rev202 version)^16^.

### sDeep homology-based protein clustering

For computing gene family clusters, we used MMseqs2^18^ with relaxed thresholds: minimum percentage of amino acids identity of 30%, E-value < =1e-3, and minimum sequence coverage of 50%. The parameters used were *--min-seq-id 0*.*3 -c 0*.*5 --cov-mode 1 --cluster-mode 2 -e 0*.*001*. We discarded clusters with less than 3 sequences. We computed Multiple Sequence Alignments for each gene family with Clustal Omega^45^ using the translated version of the genes; and subsequently reconstructed their phylogeny with FastTree2^46^. We calculated alignment statistics (mean identity, most unrelated pair, most distant sequence) on each protein family alignment using Alistat^47^.

### Detection of protein clusters specific from uncultivated taxa

To identify genes/proteins without homologs in current genomic databases, we mapped the members of each protein family cluster against: **i)** EggNOG v5^24^ with eggNOG-mapper v2^48^ searches on all the protein sequences of each family. Hits with an E-value < 1e-3 were considered significant; **ii)** PFamA^49^ with HMMER^50^ hmmsearch against all the protein sequences of each family. Hits with an E-value < 1e-5 were considered significant. **iii)** PFamB^23^ with HMMER hmmsearch searches against the representative protein sequences (longest sequences) of each family. Hits with an E-value < 1e-5 were considered significant. **iv)** Refseq^51^ with DIAMOND^52^ blastx (sensitive flag) searches against the cds sequences of all the members of the family. Hits with an E-value < 1e-3 and query coverage > 50% were considered significant. All protein family clusters with a significant hit in any of the above databases were considered non-novel and discarded from the study.

### Detection of spurious domains in protein families

In order to discard sequencing errors and potential pseudogenes, we mapped our catalog against the AntiFAM ^26^ database (HMMER search, *--cut_ga parameter*, as recommended), and discarded families with E-value < 1e-5. We also discarded families with significant hits in the viral PVOG database^27^ using HMMER and an E-value threshold < 1e-5 and minimum coverage of 50%.

### Conserved Domain Detection

We generated Multiple Sequence Alignments for each family with Clustal Omega. For each protein family, the most conserved domain was considered the longest aligned region in which 80% of the residues were not gaps. Protein families whose most conserved domain was shorter than 20 residues were discarded.

### Calculation of dN/dS

Multiple sequence alignments from each protein family were back-translated into codon alignments, and used to reconstruct phylogenetic trees using FastTree2 with default parameters. The whole workflow was executed using ETE v3^53^ with options *ete3 build --nt-switch-threshold 0*.*0 --noimg --clearall --nochecks -w clustalo_default-none-none-none --no-seq-rename*. To calculate selective pressure per family, we ran HyPhy with the BUSTED model ^54^ and default parameters using the nucleotide alignment and tree generated previously, and retrieved the dN/dS under the full codon model. We discarded protein families with dN/dS values higher than 0.5.

### Detection of protein coding families

We used back-translated alignments for running RNAcode^55^. Because the software calculates statistics on the longest sequence, we rearranged the alignments so that the longest sequence was the first to appear. We ran RNAcode with default options and the *--stop-early* flag. Protein families yielding RNAcode p-values lower than 0.05 were considered coding and retained in our catalog. Protein families without significant p-value were discarded from the study, unless they were detected in metatranscriptomics datasets (see next section).

### Mapping to metatranscriptomic datasets

In order to obtain additional evidence of gene expression, we mapped our protein families against i) TARA oceanic metatranscriptomic catalog^2^ and ii) 756 human gut metatranscriptomic samples^56^. For mapping the sequences of the novel families against TARA v2 protein sequences we used DIAMOND blastp with the *sensitive* flag. We considered any hit with E-value < 1e-3 and query coverage > 50% as significant. For mapping reads from the human gut metatranscriptome samples against the sequences of the novel families, we used DIAMOND blastx sensitive mode. We considered any hit with E-value < 1e-3 as significant. We considered families to be expressible if at least one member had a significant match against the metatranscriptomic catalogs.

### Orthology calling

Protein family clusters were analyzed in order to determine whether they represent basal orthologous groups at the bacterial or archaeal level, or, by contrast, contained duplication events leading to several orthologous groups within the same family. To do so, we rooted the phylogenetic tree of each family at midpoint and taxonomically annotated leaf nodes using GTDB v202. Then, we used ETE^53^ to identify duplication events in the tree topology of each family and date them according to their predicted common ancestry.

### Comparisons with eggNOG

We compared the distribution of protein lengths in eggNOG v5 orthologous groups^24^ and the small peptides catalog described in^12^ with the length of the novel protein families. The length of each family or orthologous group was set to the longest protein sequence within each cluster.

### Sequence based predictions

We ran SignalP-5.0^57^ with *gram +* and *gram -* modes on the protein families. Genes predicted to have a signal peptide by the *gram +, gram -* or both modes were considered as secreted proteins. We also ran TMHMM^58^ with default parameters to calculate transmembrane domains on the sequences. A protein family was considered transmembrane / secreted if at least 80% of the members of the family were predicted to be so. The same procedure was followed for measuring the proportion of transmembrane and signal peptide families on bacterial and archaeal eggNOG groups.

### Small peptide predictions

We considered families whose longest sequence was shorter than 50 residues to be small peptides. We mapped our set of small peptides against those described by Sberro *et al*., using DIAMOND (*sensitive* flag), and considered as significant those with E-value < 1e-3 and coverage > 50%. We ran antimicrobial predictions on our small peptide novel families using Macrel^59^.

### Detection of mobile elements

For detecting families potentially included in plasmids, we ran PlasFlow^60^ with the *--threshold 0*.*95* flag on all the contigs from the 5 MAG collections. For detecting families with potential viral origin, we ran Seeker ^61^ on all the contigs from the 5 MAGs datasets, using a 0.9 threshold for considering a sequence as viral. For results shown in Figure 3, we considered protein families as mobile or viral if at least one member of the family was predicted so. The reported correlations are also present under more restrictive thresholds where at least 30% of the protein family members were predicted as mobile or viral sequences (Figure S4).

### Computation of ecological distribution

For expanding the ecological profile of the novel protein families, we mapped a representative sequence member of each family against 63,410 public metagenomic samples using DIAMOND (-*sensitive* flag). Hits with an E-value lower than 10E–3 and target coverage ≥ 50% were considered as significant. We used the beta_diversity package from scikit-bio with *braycurtis* mode to calculate the beta diversity of the samples in which each family was detected.

### Taxonomic breadth of protein families

For estimating the taxonomic breadth of each novel protein family, we calculated the Last Common Ancestor (LCA) of their members using GTDB v202 taxonomic predictions as the most lineage specific annotation shared by all members of the family. To make sure that our LCA predictions were not artifacts caused by a small proportion of missanotated genes masking lineage specific families to very basal levels, we repeated the analysis requiring the LCA lineage to be supported by 80% (Figure S5) and 50% (Figure S6) of the members of the family, obtaining comparable patterns.

### Taxonomic distribution of novel protein families

To assess the distribution of novelty across the prokaryotic phylogeny, we estimated the amount of novel protein families observed per clade. Figure 2 shows the GTDB bac120_r202 and ar122_r202 phylogenetic trees collapsed to the order level (each leaf represents a taxonomic order). For representing the percentage of uncultivated genomes per branch in Figure 2, we divided the number of uncultivated genomes per lineage by the total number of genomes under that lineage. The final tree image was generated using iTOL^62^.

### Synapomorphic protein families

For identifying synapomorphic protein families at different taxonomic levels, we calculated the clade specificity and coverage of each protein family across all GTDB v202 lineages. For each protein family and clade, coverage was calculated as the number of genomes containing a specific protein family over the total number of genomes under the target clade. Specificity was estimated as the percentage of protein members within a family that belonged to the target clade. We considered protein families as synapomorphic if they contained at least 10 members (i.e., protein sequences from different genomes) and had a coverage higher than 0.9 and a specificity of 1.0 for a given lineage. Moreover, to further ensure that our synapomorphic predictions were highly specific, we mapped them back against the whole catalog using a more sensitive mapping strategy based on HMMER searches^50^ and excluded families with distant hits to genomes that might compromise the strict specificity and coverage thresholds of 1.0 and 0.9 respectively.

### Functional predictions

In order to infer the functional annotation of the neighboring genes of the novel families, we ran eggNOG-mapper v2^48^ with default parameters on the 400M proteins in our catalog of MAG, SAGs and reference genomes. We also mapped them against the CARD^63^ database for retrieving their functional annotations, using diamond blastp with e-value and coverage threshold of 10E–3 and > 50% respectively.

For the 5 MAG sets, we built a database with all the neighbor genes and their positions in each scaffold. We next measured the functional conservation in a genomic window of +-3 genes around each targeted novel gene. For this, we calculated the prevalence of functional terms (eggNOG orthologous group, KEGG pathway, KEGG orthology, KEGG module, PFAM, CAZy and CARD) along the neighboring genes of all members within the same protein family. We obtained a conservation score for each observed functional term as: *Conservation (position X, annotation Y) = number of genes in position X with annotation Y / number of members in the family*. We also took into consideration whether the neighboring genes were in opposite directions than the novel gene as well as the distance between genes. Predictions for novel protein families were inferred if the conservation score for that pathway was higher than 0.9 and the neighboring genes used as functional-term donors were located in the same DNA strand as the unknown genes and at a maximum distance of 100 nt. We followed the same procedure for finding families consistently located next to resistance genes in the CARD database.

For locating families putatively involved in relevant processes, we manually selected KEGG orthologous groups (KOs) that were highly specific for metabolic pathways involved in energy metabolism and xenobiotic degradation. In particular, we selected KOs involved in a maximum of two pathways, therefore discarding most promiscuous molecular functions and considering only specific biomarkers. The list of selected KOs is provided in Table S7. To create figure 4, we joined the bacterial and archaeal GTDB phylogenetic trees collapsed to the order level, using the ETE toolkit software. Then, we represented the presence/absence matrix of novel protein family predictions associated with pathway specific KO across the different taxonomic orders. For readability, orders with no predictions are not shown in the figure.

We also mapped the novel family sequences against the Fitness Browser^33^ genes (https://fit.genomics.lbl.gov/cgi_data/aaseqs) using DIAMOND-blastp with e-value and coverage threshold of 10E–3 and > 50% respectively. We considered hits with strong fitness changes (t-score > 4 or < -4, see original publication) to be potentially associated with certain conditions.

## Data and material availability

The genomic context, sequence signatures and taxonomic distribution of each gene family can be visualized at http://novelfams.cgmlab.org/. This site includes a Downloads section from where the raw sequences, alignments and hmm files of all families can be downloaded. The code used for generating these results and the supplementary tables were uploaded to https://github.com/AlvaroRodriguezDelRio/NovFamilies.

## Acknowledgments

We thank Yiqian Duan and Célio Dias Santos Júnior (Fudan University) for their help with the habitat annotation of metagenomes.

## Supplementary Tables

All supplementary tables are provided within the following Excel file: https://github.com/AlvaroRodriguezDelRio/NovFamilies/raw/main/Supplementary%20tables.xlsx

## Supplementary Figures

**Figure S1.**
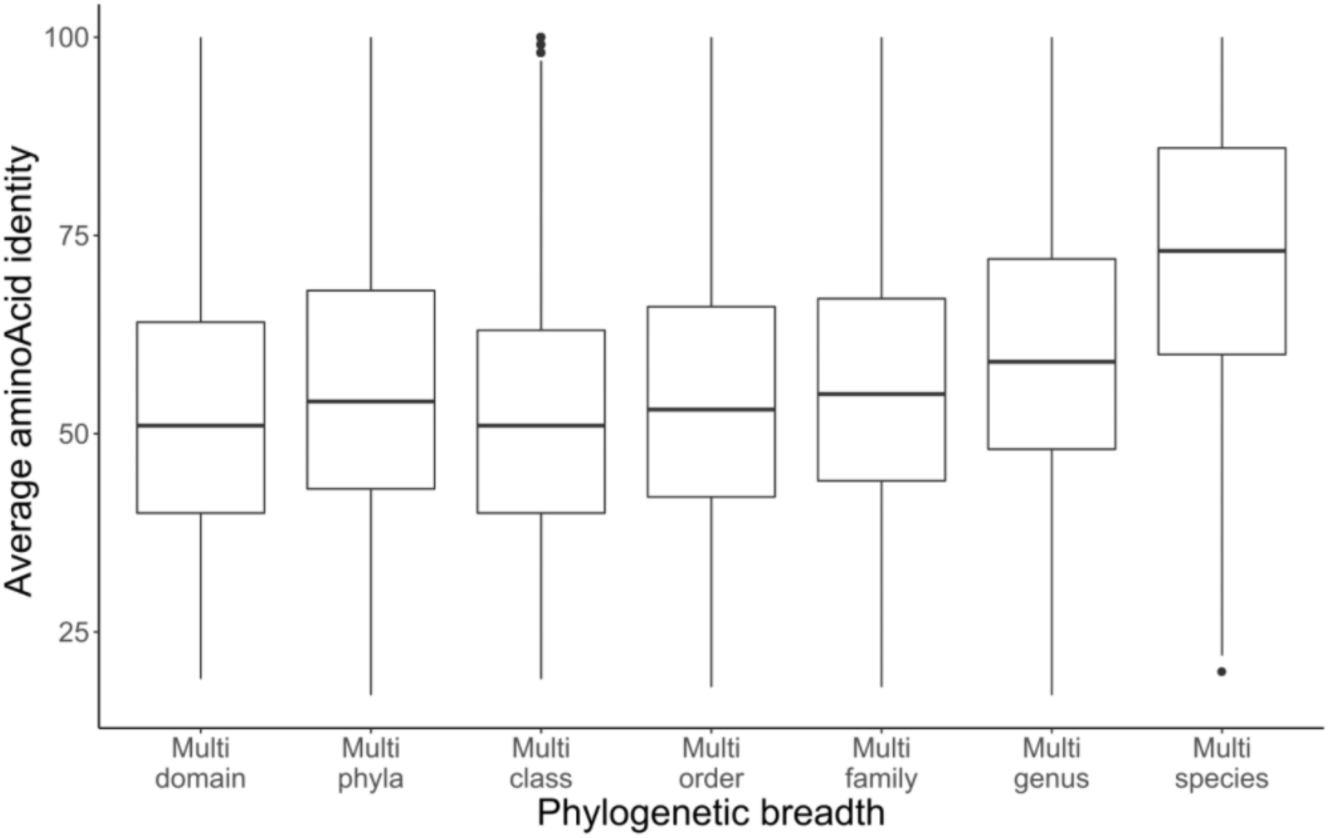
Average amino acid identity of the families, stratified by taxonomic breadth.

**Figure S2.**
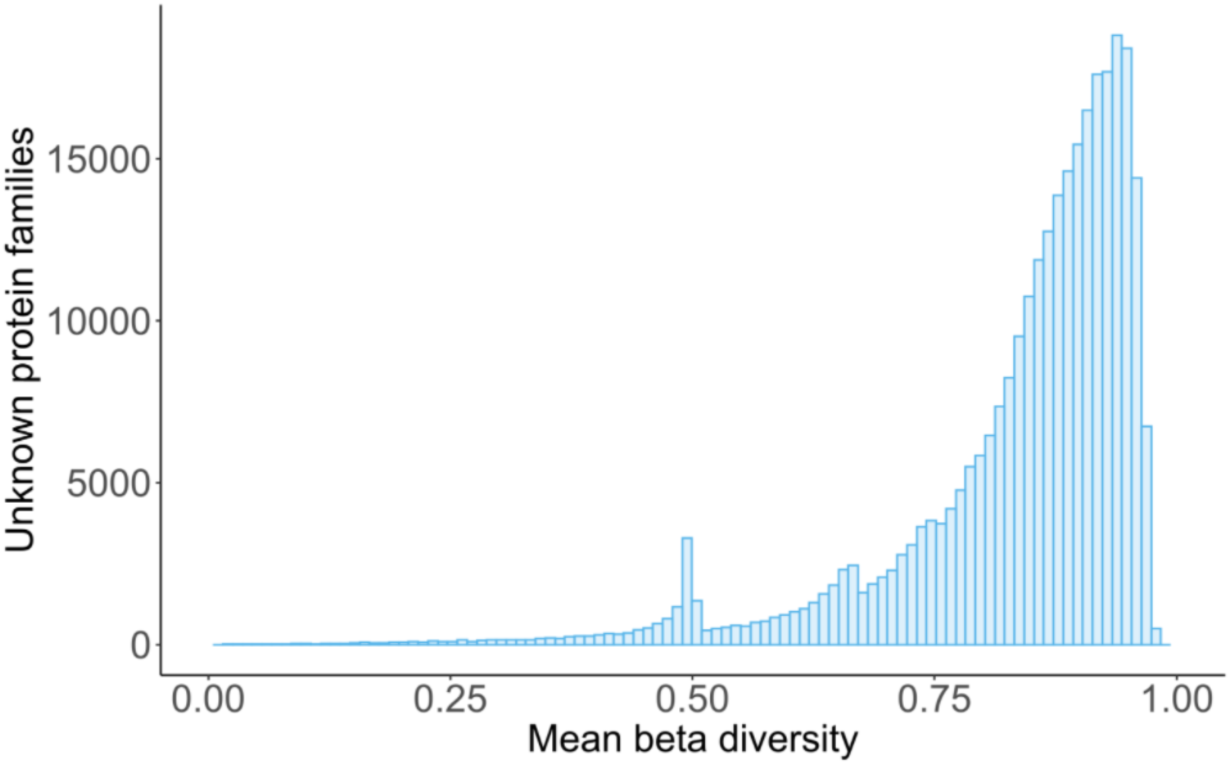
Distribution of beta diversity values calculated for the novel families.

**Figure S3.**
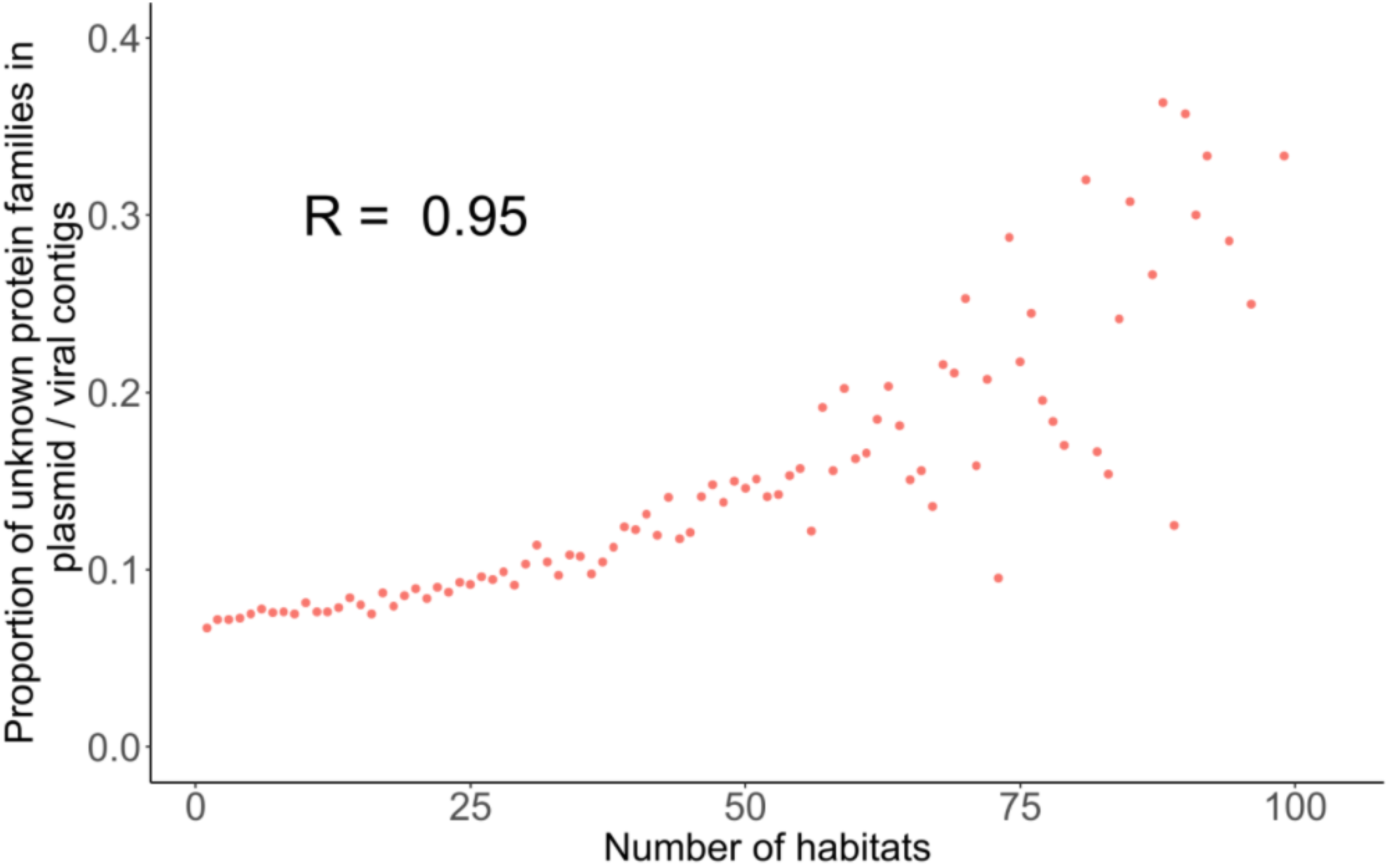
Proportion of protein families linked to plasmids or viral contigs with relation to the number of habitats they were detected in. R was calculated as the Spearman correlation coefficient.

**Figure S4.**
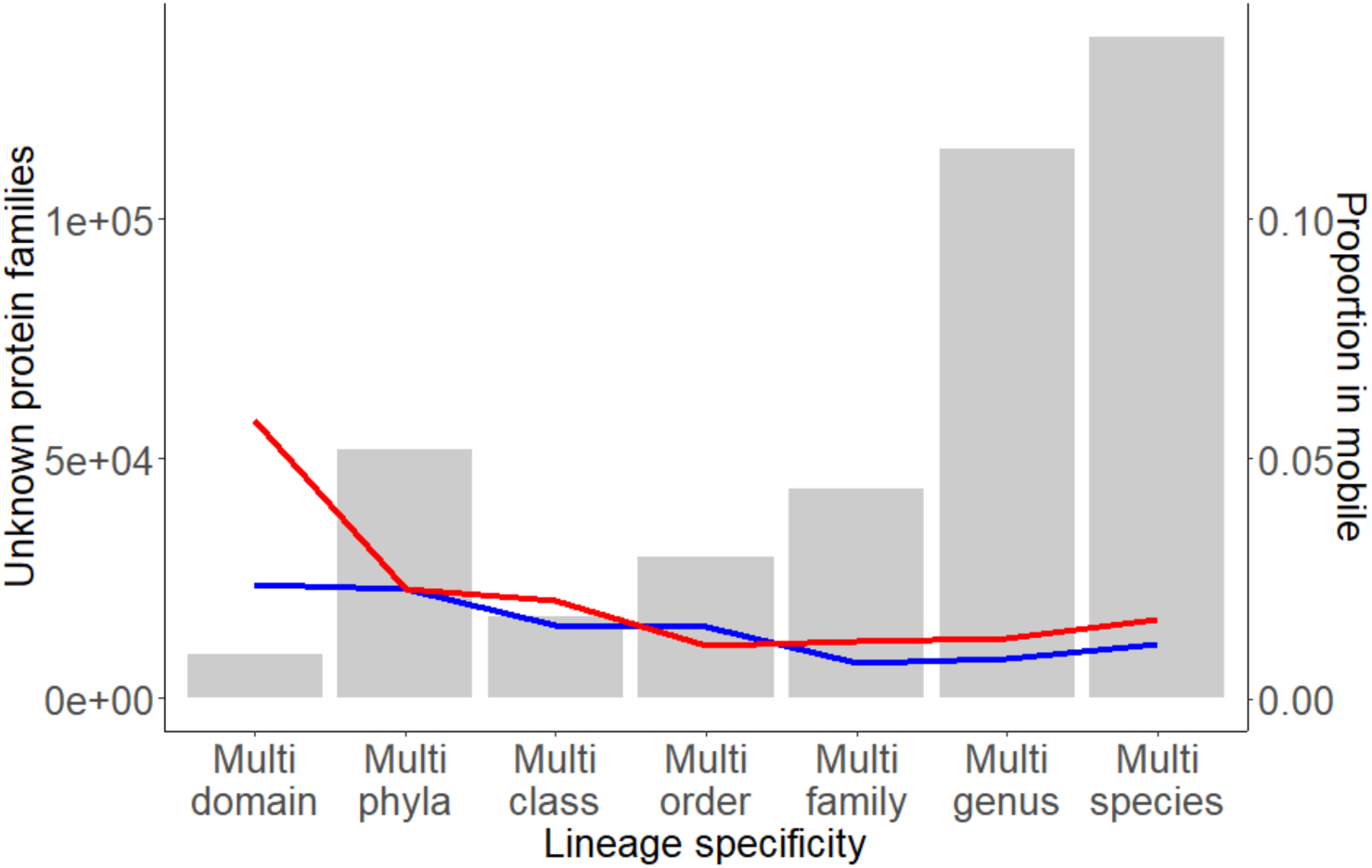
Equivalent to Fig. 3C but requiring 30% of the members to be present in plasmids (blue) / viral contigs (red) for considering the family as mobile.

**Figure S5.**
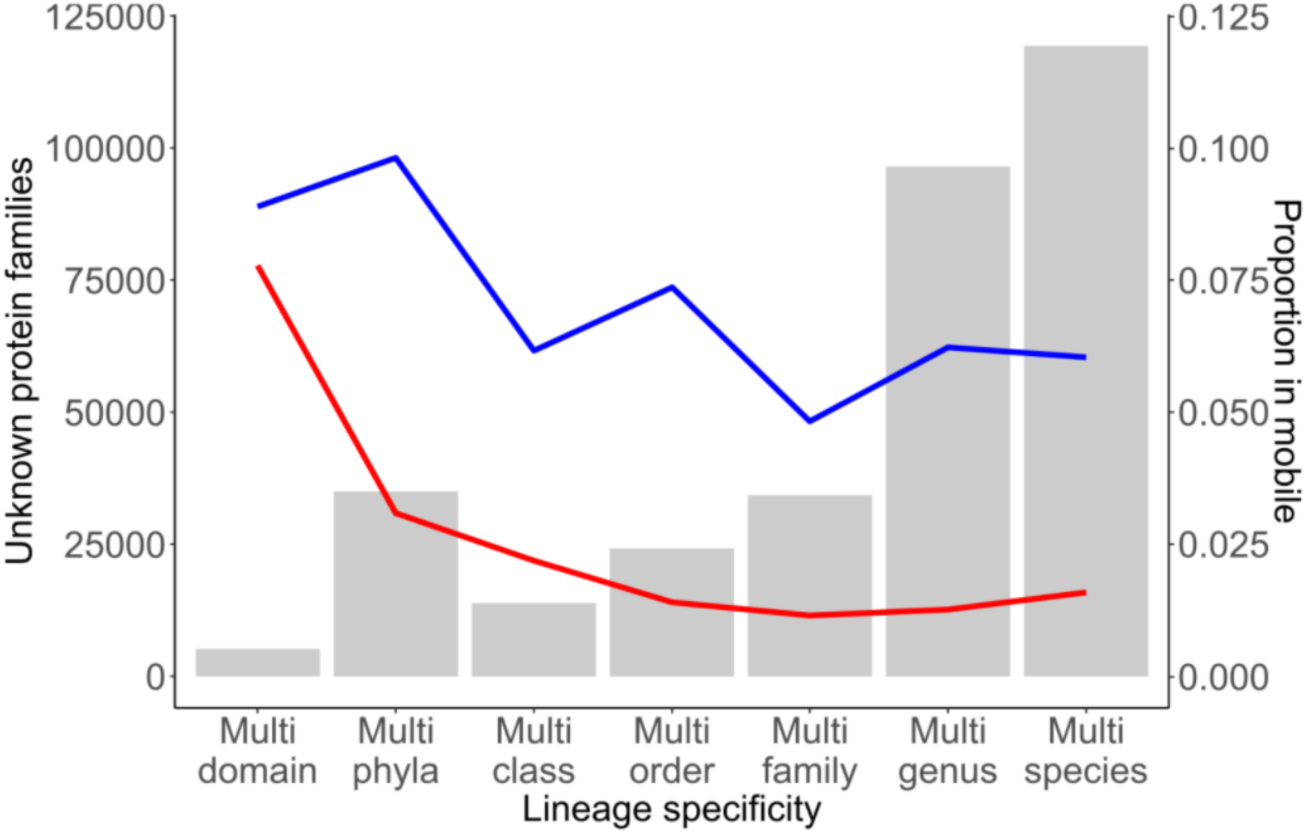
Equivalent to Fig. 3B but calculating LCAs as the most basal taxonomic group gathering 80% of the members of the family. Red: proportion of families in viral contigs. Blue: proportion of families in plasmids.

**Figure S6.**
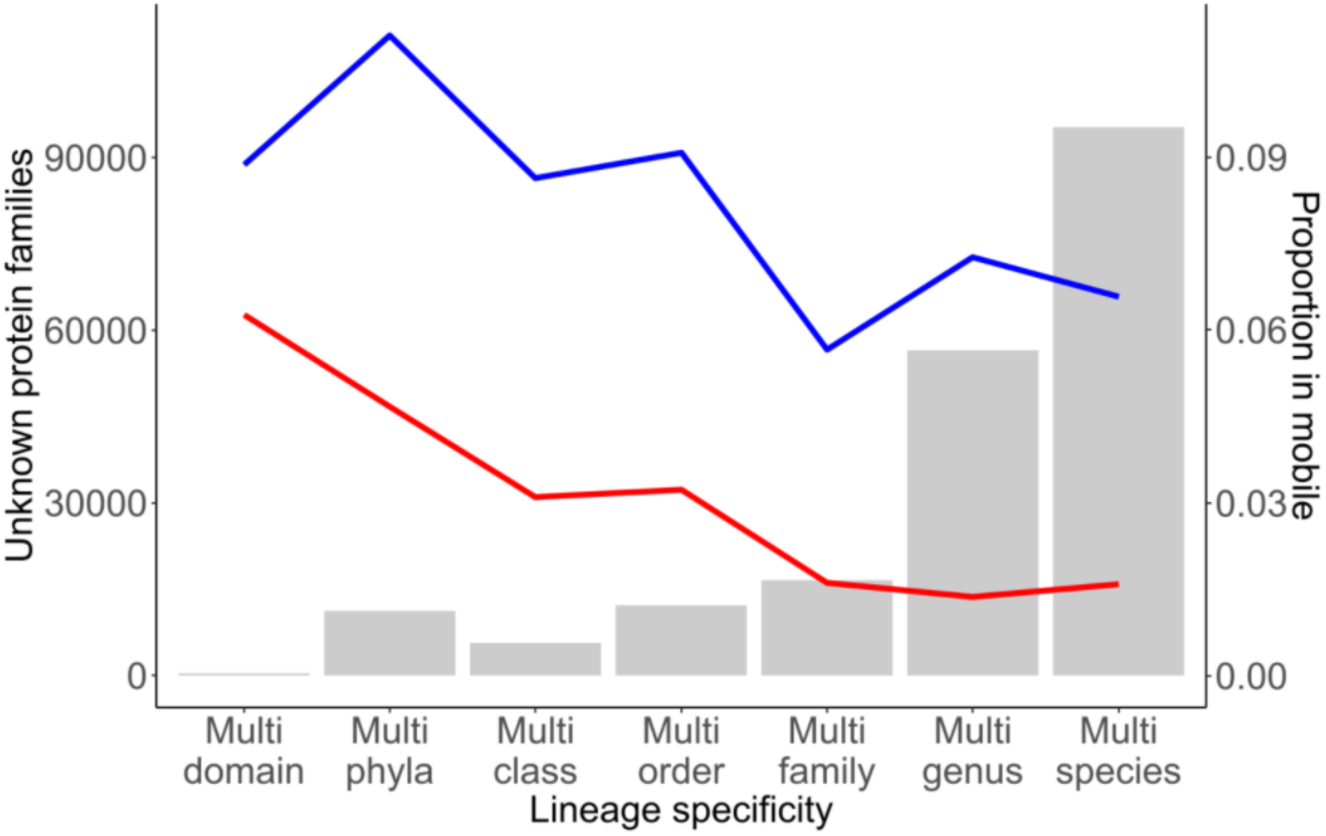
Equivalent to Fig. 3B but calculating LCAs as the most basal taxonomic group gathering 50% of the members of the family. Red: proportion of families in viral contigs. Blue: proportion of families in plasmids.

